# Diet Induced Obesity and Diabetes Enhance Mortality and Reduces Vaccine Efficacy for SARS-CoV-2

**DOI:** 10.1101/2022.10.15.512291

**Authors:** Robert M Johnson, Jeremy Ardanuy, Holly Hammond, James Logue, Lian Jackson, Lauren Baracco, Marisa McGrath, Carly Dillen, Nita Patel, Gale Smith, Matthew Frieman

**Affiliations:** Center for Pathogen Research, Department of Microbiology and Immunology, University of Maryland School of Medicine, Baltimore, MD, 20201; Novavax, Gaithersburg, MD

## Abstract

Severe Acute Respiratory Syndrome Coronavirus-2 (SARS-CoV-2), the causative agent of Coronavirus disease 2019 (COVID-19), emerged in Wuhan, China, in December 2019. As of October 2022, there have been over 625 million confirmed cases of COVID-19, including over 6.5 million deaths. Epidemiological studies have indicated that comorbidities of obesity and diabetes mellitus are associated with increased morbidity and mortality following SARS-CoV-2 infection. We determined how the comorbidities of obesity and diabetes affect morbidity and mortality following SARS-CoV-2 infection in unvaccinated and adjuvanted spike nanoparticle (NVX-CoV2373) vaccinated mice. We find that obese/diabetic mice infected with SARS-CoV-2 have increased morbidity and mortality compared to age matched normal mice. Mice fed a high-fat diet (HFD) then vaccinated with NVX-CoV2373 produce equivalent neutralizing antibody titers to those fed a normal diet (ND). However, the HFD mice have reduced viral clearance early in infection. Analysis of the inflammatory immune response in HFD mice demonstrates a recruitment of neutrophils that was correlated with increased mortality and reduced clearance of the virus. Depletion of neutrophils in diabetic/obese vaccinated mice reduced disease severity and protected mice from lethality. This model recapitulates the increased disease severity associated with obesity and diabetes in humans with COVID-19 and is an important comorbidity to study with increasing obesity and diabetes across the world.

**Importance:** SARS-CoV-2 has caused a wide spectrum of disease in the human population, from asymptomatic infections to death. It is important to study the host differences that may alter the pathogenesis of this virus. One clinical finding in COVID19 patients, is that people with obesity or diabetes are at increased risk of severe illness from SARS-CoV-2 infection. We used a high fat diet model in mice to study the effects of obesity and Type 2 diabetes on SARS-CoV-2 infection as well as how these comorbidities alter the response to vaccination. We find that diabetic/obese mice have increased disease after SARS-CoV-2 infection and they have slower clearance of virus. We find that the lungs of these mice have increased neutrophils and that removing these neutrophils protect diabetic/obese mice from disease. This demonstrates why these diseases have increased risk of severe disease and suggests specific interventions upon infection.

## Introduction

The United States of America has an obesity rate of 42%, diabetes rate of 11%, and 35 % of the population has been identified as prediabetic[1]. Diabetes and obesity have been linked both clinically and through physiological studies to increased severity of many types of diseases that can be traced toward various aspects of these metabolic syndromes[2-4]. Epidemiologic studies of COVID-19 have identified obesity and diabetes as significant risk factors for increased disease severity, length of hospitalization, and death[5-10]. The immune and molecular mechanisms behind this increased risk of COVID-19 have yet to be fully determined. SARS-CoV-2 is not the only coronavirus that displayed increased disease severity in those with diabetes or obesity [6, 11-17]. The Middle East respiratory syndrome coronavirus (MERS-CoV), first identified in 2012, was clinically and experimentally linked to increased disease severity in those with diabetes[18, 19]. Across multiple clinical and epidemiological studies, diabetics have a 3-7-fold higher odds ratio of severe disease and death after MERS-CoV infection[11, 14-16]. Mouse models of disease and viral infections have been used for studying coronaviruses and other pathogens[20]. MERS-CoV also demonstrated increased morbidity and mortality in an obese/diabetic mouse model, with high fat diet (HFD) mice presenting with increased lung pathology, viral titer, and a dysregulated immune response to MERS-CoV infection [11, 19, 21].

The etiology of type 2 diabetes is closely linked with chronic inflammation induced by excess adipose tissue. Stressed adipocytes and adipose tissue macrophages (ATMs) secrete numerous proinflammatory cytokines that result in chronic low-grade inflammation which alters homeostatic glucose regulation by decreasing cellular responsiveness to insulin[4]. This results in hyperglycemia, hyperinsulinemia, and glucose intolerance characteristic of type 2 diabetes (T2D). The systemic inflammatory responses of obesity and T2D are seen in the microenvironment of the lungs. Polarization of the macrophage response is observed where a Th1 response predominates in OB/OB and DIO mice[22, 23]. It is shown that fewer macrophages are recruited to the lungs which allows for the inflammatory response to recruit other cell types, including neutrophils which we observe in this manuscript[24, 25]. This systemic inflammatory response leads to altered vaccine responses for a variety of vaccines in humans and can be observed in mice. Through several studies, the role of T cells is dominant where vaccinated obese or diabetic mice have reduced T cell activation thus altering the vaccine efficacy[26]. The mechanism underlying these deficiencies are currently unknown.

To study the role of obesity and diabetes in SARS-CoV-2, we utilized a mouse model where mice are fed a high fat diet (60% fat from diet) starting at 3 weeks of age compared to a normal diet (ND) (22% fat from diet). By 14 weeks of age, HFD mice are obese, glucose intolerant and have high resting glucose levels; all metrics of having Type 2 diabetes (T2D). Infection of ND and HFD mice with SARS-CoV-2 (MA10 mouse adapted strain) demonstrates lethal disease in HFD only mice by 7 dpi. Equivalent viral lung titers were found in both ND and HFD mice, but increased lung inflammation was observed in HFD (obese/diabetic) mice. Vaccination of ND and HFD mice with the saponin-based Matrix-M adjuvanted spike nanoparticle (NVX-CoV2373)[27, 28] resulted in generation of similar levels of neutralizing antibody. However, when challenged with SARS-CoV-2 (MA10), HFD mice had a delay in clearance with little reduction in lung titer. Evaluation of the inflammatory response in HFD vaccinated/infected mice found an increase in neutrophils in lung infiltrate that correlated with an increased innate inflammatory response. When neutrophils are depleted in HFD mice we observed reduced weight loss, reduced clinical signs of disease and increased survival demonstrating a role for neutrophils in severity of disease in diabetic/obese mice. We conclude this demonstrates a dysregulated immune response in obese/diabetic mice leading to increased morbidity and mortality as well as potentially skewing the quality of protection from vaccination. This supports what is seen in diabetic and obese humans throughout the COVID-19 pandemic and provides a model to identify the molecular and immunological mechanisms underlying the response to SARS-CoV-2 in this population.

## Results

### High fat diet fed mice have obesity and type 2 diabetes

To study the effects of obesity and diabetes on SARS-CoV-2 infection, we established a model of SARS-CoV-2 infection based on the previously established HFD fed model. At the time of weaning (∼3 weeks), C57BL/6 mice are placed on a ND (18% kcal from fat, 24% kcal from protein, and 58% kcal from carbohydrates) or a HFD (60% kcal from fat, 18% kcal from protein, and 21% from carbohydrates) (Figure 1A). At 18 weeks of age, 15 weeks on each diet, we analyzed fasting blood glucose (Figure 1B), weight (Figure 1C) and *ACE2* expression (Figure 1D). The HFD mice had increased fasting blood glucose and higher weight, which are all parameters of obesity, insulin resistance and T2D. We examined the expression of *ACE2* to ensure there were no intrinsic differences that could confound results during infection. We did not observe any difference in ACE2 levels in normal vs obese/diabetic mouse cohorts. Once established, 18-week-old ND and HFD mice were then used for pathogenesis and vaccination experiments throughout the study.

**Figure 1:**
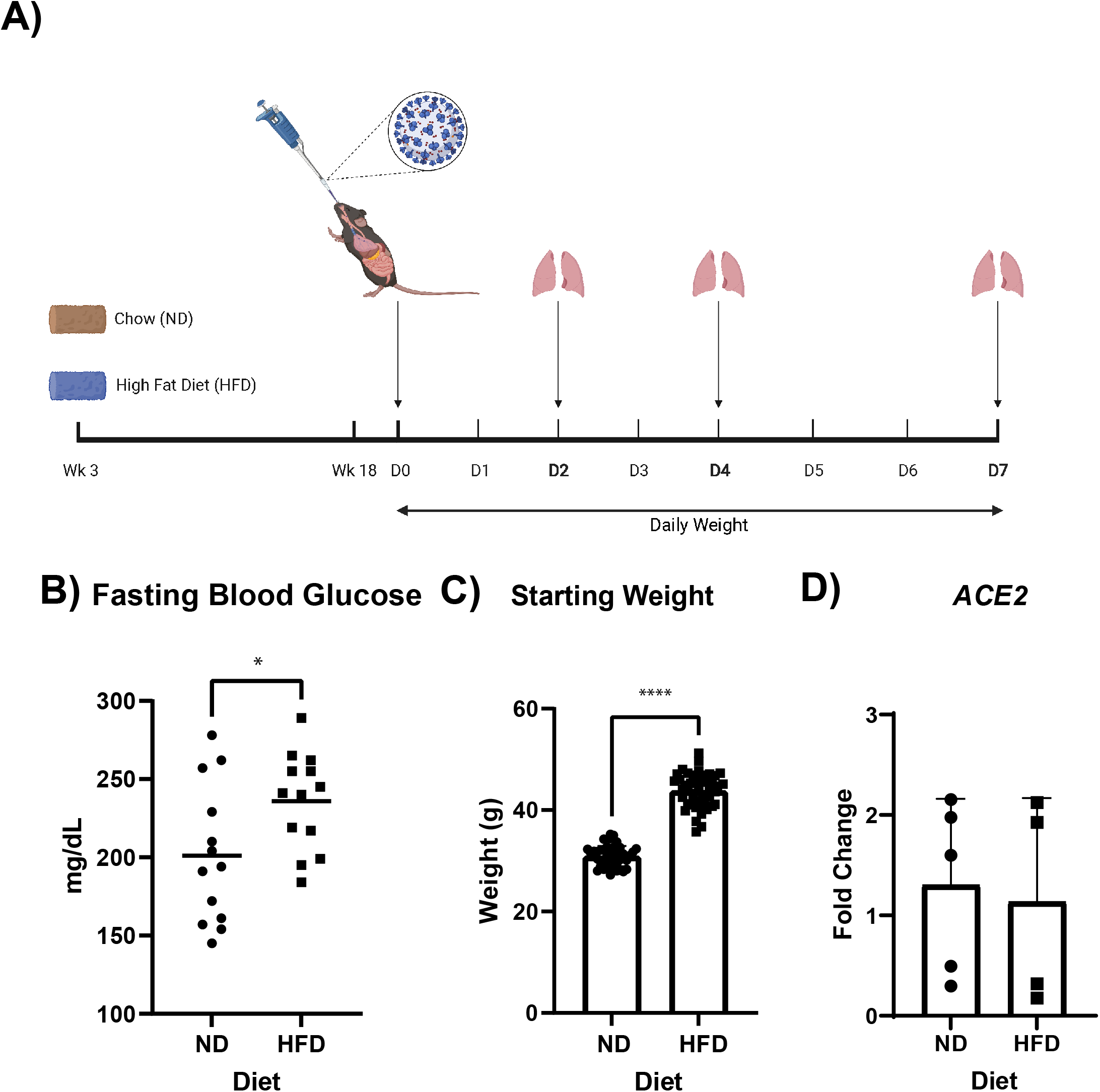
A high fat diet induces obesity and diabetes in C57BL/6J mice. **A**. Schematic overview illustrating diet schedule and infection for control and obese/diabetic mice. Briefly, at weaning (3 weeks), mice were placed on a normal diet (ND) or high fat diet (HFD). At 18 weeks of age, these mice were infected with either 1×10^3^ or 1×10^5^ pfu of SARS-CoV-2 Mouse Adaptive 10, monitored daily for weight changes, and a subset was euthanized at day 2, 4, and 7 to analyze lung viral titer and host immune response. The **B**. fasted blood glucose concentration *n=13* mice/group, **C**. The starting weight of ND and HFD mice *n=45* mice/group, and **D**. *ACE2* expression of ND and HFD mice *n=5* mice/group prior to infection. * p <0.5 and **** p < 0.0001 as determined by an unpaired two-sided t-test. The data are presented as mean ± SD.

### SARS-CoV-2 infected obese/diabetic mice have increased morbidity and mortality

ND and HFD mice were infected with the SARS-CoV-2 (MA10) strain [29] at 10^3^ and 10^5^ PFU per mouse. Mice were weighed daily for 7 days, with lungs evaluated on 2-, 4- and 7-days post-infection (dpi). ∼20% of the ND mice infected with 10^3^ or 10^5^ PFU MA10 died by 7 dpi compared to the HFD mice infected with 10^3^ PFU where ∼40% die by day 7 (Figure 2A). Most notably, 100% death was observed in the 10^5^ PFU infected HFD group. The kinetics of lethality varied between groups as well, with 20% of the HFD mice dying by day 3 whereas the ND mice did not die until 5 dpi.

**Figure 2:**
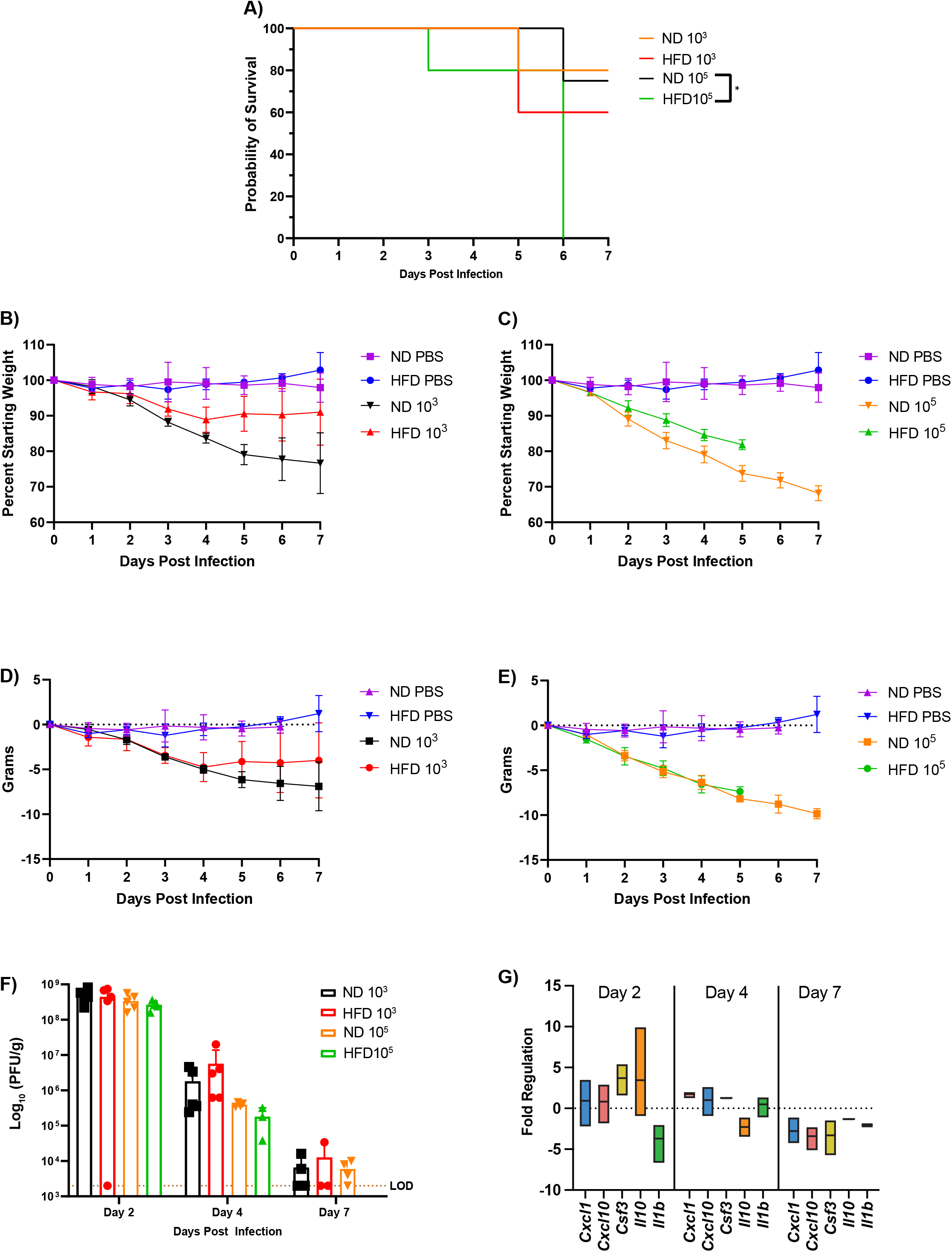
Enhanced mortality in obesity/ diabetes mice after SARS-CoV-2 infection. The survival (A), percent weight loss for low dose (B), high dose (C), grams lost for low dose (D), and high dose (E) challenge. Viral lung titers were assessed for control and obese/ diabetic mice throughout a SARS-CoV-2 MA10 infection (F). For **A**. * p < 0.05 as determined by a Log-rank (Mantel-Cox) test for survival analysis between ND and HFD for low and high infection dose with the data presented as mean ± SD with *n=10* mice/group. In G, lung cytokine, chemokine, and growth factors were assessed at 2, 4, and 7 dpi for the 1×10^3^ infectious dose using a mouse cytokine/chemokine array. The fold change was calculated compared to ND infected mice (n=3). For **B-D** the data are presented as mean ± SD with *n*=3-5 mice/group.

Weight loss was followed through 7 dpi in all groups (Figure 2B-E). For the low infection dose, we observed the ND mice starting to lose weight at 2 dpi with a total weight loss of 20% by 7 dpi. The HFD mice lost weight starting 2 dpi, which continued until 4dpi, for a total loss of around 10% and was maintained for the remainder of the experiment. For the high infection dose, the ND mice lost weight starting on 2 dpi and continued through the day 7 endpoint of this experiment with a total weight loss of ∼30% (Figure 2C). The HFD mice lost weight starting on 2 dpi and continued until succumbing to the infection on 5 dpi (Figure 2C). As ND mice weigh less than HFD mice, analysis of by grams of weight lost intra group, as compared to total %, can provide additional insight. Here, when we compare the ND and NFD mice, no differences were observed (Figure 2D & E).

We next examined viral replication in the lungs of infected mice at 2, 4, and 7 dpi. For all groups, the peak viral replication occurred at 2 dpi with a viral titer ∼10^8^ pfu/g of lung and decreased through day 7 (Figure 2F). At 4 dpi, we observed less viral titer in the 10^5^ PFU group compared to the 10^3^ PFU group (Figure 2F). This was surprising; however, we suspect that the reduced viral titer in the high dose lungs is potentially due to either reduced numbers of infectable cells at this timepoint due to virus induced cell death or a large influx of inflammatory cells, potentially limiting replication but not to substantial levels or quality to protect from death.

### Obese/diabetic mice have altered host immune responses after infection with SARS-CoV-2

As obesity/diabetes is known to augment the host immune response[30-34], we examined the cytokine and chemokine profile of uninfected, SARS-CoV-2 infected ND and HFD mice to determine if differences in immune response correlated with disease. To perform a longitudinal analysis, we used the samples from mice infected with 1×10^3^ pfu/mouse across 7 dpi. qPCR was performed on lung RNA, and we investigated the changes in expression for a variety of chemokine and cytokine genes in both ND and HFD mice. On days 2 and 4 post-infection, we observed a neutrophil recruitment associated chemokines in HFD compared with ND mice (Figure 2G). Cxcl1, Cxcl10, Csf3 and IL10 are all key neutrophil recruitment factors and are highly upregulated in HFD mice at day 2 of infection. Those levels are reduced at day 4 and then lower at day 7. This correlates with the neutrophil infiltration kinetics seen in the lungs as shown in Figure 4. Interestingly, we observe that HFD mice had a ∼5 fold down-regulation of IL1-β, which is a mediator of lung inflammation, fever and fibrosis and is associated with protection in human COVID19 patients (Figure 2G). We hypothesize that neutrophils play an early and critical role in protection from lethal disease in this model and that reduction of neutrophils may protect mice from lethal disease.

### Obese/diabetic mice have no difference in lung pathology after infection with SARS-CoV-2

At the low infectious dose, both HFD and ND mice show similar lung pathology characterized by a mixed inflammatory infiltrate that peaks at 4 dpi and begins to resolve by 7 dpi. Epithelial cell sloughing in the bronchi is observed starting at 2 dpi with epithelial cells replaced by 7 dpi in most large airways. An observable level of edema is also visible in both HFD and ND lungs across time points (Figure 3A).

**Figure 3:**
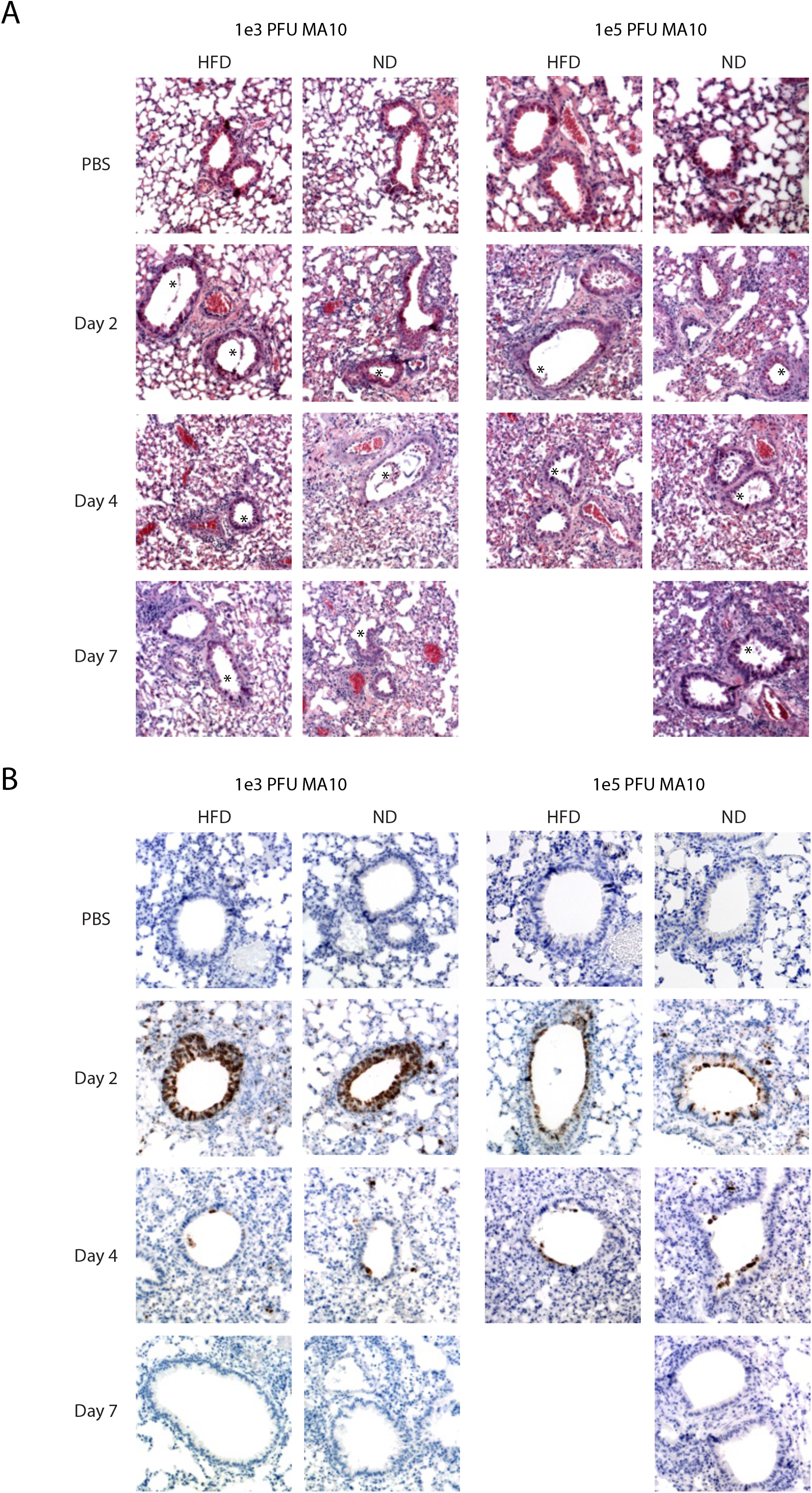
Similar lung histology and viral clearance in control and obese/ diabetic mice. Control and obese/diabetic mice were infected intranasally with either 1×10^3^ or 1×10^5^ pfu/ mouse. Lungs were collected on 2, 4, and 7 dpi and fixed in 10% neutral buffered formalin for greater than 24 hours. Tissue was embedded in paraffin. **A**. 5 µm sections were cut and stained with hematoxylin and eosin, and **B**. immunostained with anti-SARS Nucleocapsid Protein antibody. Images are shown at original magnification x10 and are representative of *n* = 3-5 mice/group.

At the high infectious dose, the course of inflammation in the lung is similar but more severe than at the low dose. Mixed inflammatory infiltrates in the alveolar space are observed beginning at 2 dpi and increasing through 7 dpi for the surviving ND mice. The infiltrates are predominantly comprised of neutrophils and monocytes, as observed by H&E and quantified by flow cytometry analysis in Figure 7. Bronchiolar sloughing is also more pronounced at the high infectious dose, with visible damage still present at 7 dpi. Perivascular cuffing and edema were also observed in the high infectious dose mice (Figure 3A).

Lung sections were stained for the presence of nucleocapsid (N) protein using immunohistochemistry to visualize viral antigen localization within the lung sections and to determine if HFD caused broader spread of infection than ND fed mice. Similar to what was observed by viral titer, the amount of N staining is highest at 2 dpi, with the majority of N localizing in the ciliated epithelial cells of the large airways for both the control and HFD mice (Figure 3B). We detect a small amount of N positivity outside the airways, most noticeably at 4 dpi (Figure 3B). Sections from 7 dpi show little to no N staining positivity, correlating with the viral clearance pattern observed by the viral titer (Figure 3B). No differences are observed between ND and HFD mice demonstrating that HFD mice have no change in virus tropism. This data demonstrates that obesity/diabetes increases the mortality of a SARS-CoV-2 infection.

### Vaccinated HFD mice suffer from weight loss and have delayed clearance after SARS-CoV-2 infection

Obesity/diabetes decreases vaccine efficacy towards viral respiratory pathogens [30-36]. In humans receiving SARS-CoV-2 mRNA vaccination, those with obese/diabetic co-morbidity demonstrate the highest rates of severe disease and hospitalization [37-40]. Thus, to mimic the timing of vaccination in the general population, we assessed the efficacy of a clinically licensed adjuvanted protein subunit vaccine (NVX-CoV2373) vaccine in obese/diabetic mice. NVX-CoV2373 is an adjuvanted full-length Spike protein vaccine based on the SARS-CoV-2/Wuhan1 sequence that is highly effective at reducing SARS-CoV-2 infection in young ND mice[28]. At 18 weeks of age, ND and HFD mice were vaccinated with 1 µg of NVX-CoV2373 and then received a second dose 14 days later. On day 28 post-vaccination, blood was collected to evaluate neutralizing antibodies towards SARS-CoV-2 (MA10). On day 32 (∼2 weeks after 2^nd^ vaccination), mice were challenged with 1×10^5^ pfu/mouse of SARS-CoV-2 (MA10) (Figure 4A). The 1×10^5^ pfu is lethal in obese/diabetic mice by 7 dpi, as seen in Figure 2A, so for this experiment the mice were only analyzed through 4dpi. We monitored weight daily and analyzed lungs on 2 and 4 dpi to determine viral lung titer and immune responses (Figure4A). As expected, the unvaccinated mice had no detectable neutralizing antibodies against MA10 (Figure 4B). Vaccinated ND mice had ∼ 1×10^4^ neut99, as did the vaccinated HFD mice (Figure 4B), demonstrating no difference between groups. Next, we observed that non-vaccinated mice in both ND or HFD mouse groups had ∼ 50% survival rate by day 3, with no further mortality through 4dpi (Figure 4C). Unvaccinated ND mice lost 10% of their starting weight through 4dpi (Figure 4D), which was prevented with vaccination. However, vaccination in HFD mice did not protect from a ∼10% starting weight loss by 4dpi.

**Figure 4:**
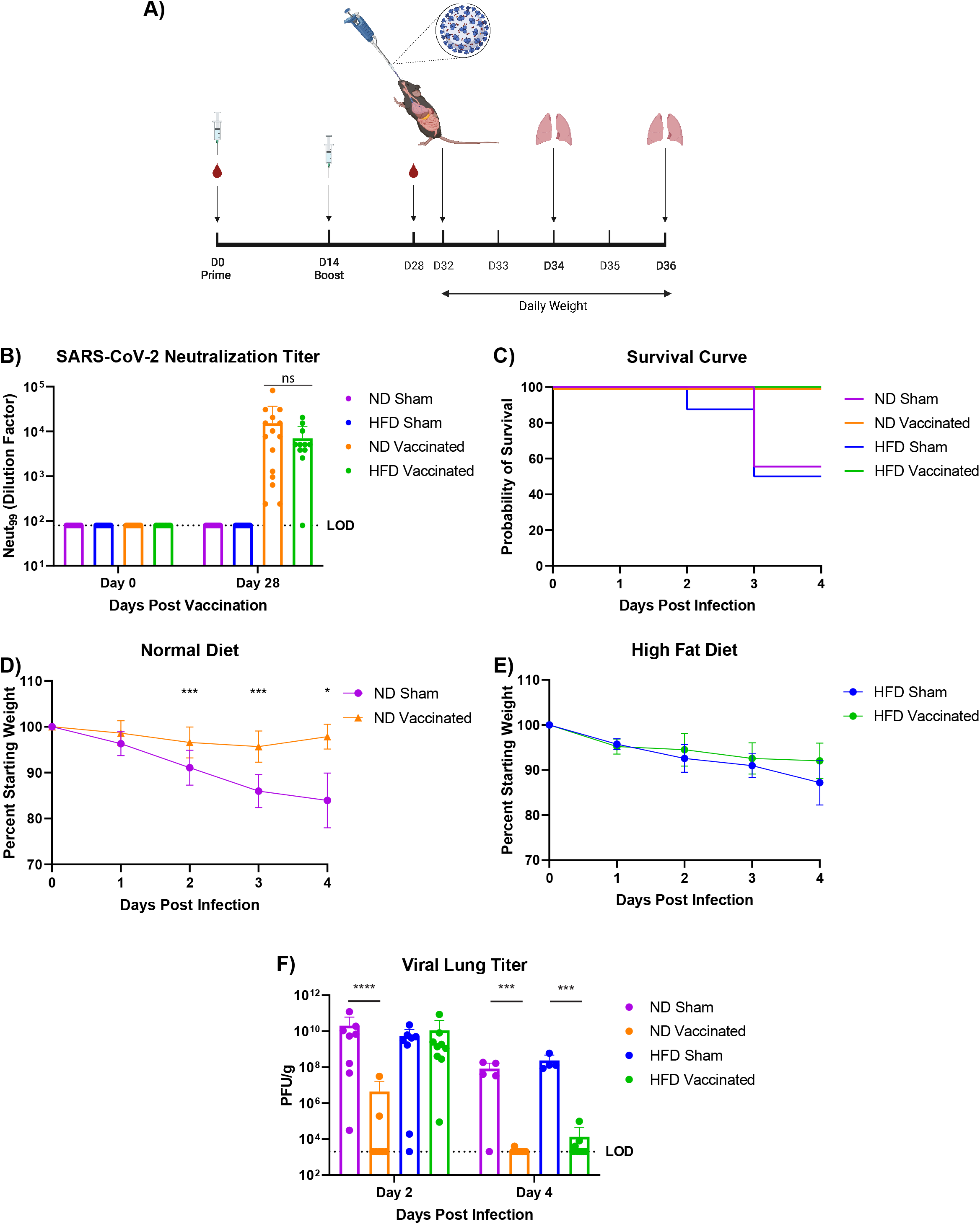
Viral clearance is delayed in obese/ diabetic vaccinated mice. **A**. Schematic overview illustrating vaccination and challenge for ND and HFD mice. Briefly, at 18 weeks of age, mice were vaccinated as well as 14 days later. On day 28 post-vaccination, mice were bled to determine **B**. serum neutralizing antibodies prior to infection, *n* = 12-15 mice/group. On day 32, these mice were challenged with 1×10^5^ pfu of SARS-CoV-2 MA 10, monitored daily for weight changes, and a subset was euthanized on day 2 and 4 to analyze lung viral titer and host immune response. The **C**. mortality, weight loss of **D**. ND, **E**. HFD, and **F**. viral titer of sham and vaccinated mice after SARS-2 MA10 challenge. *n=* 5-10 mice/group. For **D**, * p <0.5 and *** p < 0.001 as determined by a mixed-effect analysis followed by a Sidak test for multiple comparisons. For **F**. p *** < 0.001 and p **** < 0.0001, data were log-transformed and analyzed by mixed-effect analysis followed by Sidik for multiple comparisons. The data are pooled from 2 independent experiments and presented as mean ± SD.

Lungs of infected mice were next analyzed for viral titer at 2 and 4dpi. The unvaccinated ND mice had a viral titer of ∼10^10^ pfu/g (Figure 4F) and other than 2 mice with moderate levels in their lungs, cleared the virus by 2dpi. By 4dpi, the unvaccinated ND mice had 2 logs lower virus compared to 2 dpi, and the vaccinated ND mice had almost no detectable virus in their lungs (Figure 4F). For the HFD mice, at 2 dpi, there was little difference in lung titer between vaccinated and unvaccinated mice even though these mice had equivalent levels of neutralizing antibody (Figure 4F). At 4dpi, the unvaccinated HFD mice had equivalent lung titers to the ND mice, and the vaccinated HFD mice had no detectable virus in their lungs (Figure 4F). This demonstrates that despite comparable neutralizing titers in vaccinated HFD and ND mice, there is significantly delayed clearance of SARS-CoV-2 in obese/diabetic mice.

### HFD mice have increased neutrophil infiltration and higher inflammatory chemokine response after SARS-CoV-2 infection

Our data demonstrating that HFD mice suffer from increased weight loss and have delayed clearance suggests they have a distinct immune response that alters infection. To assess the innate immune response of HFD mice after challenge with SARS-CoV-2, we performed both flow cytometry based immunophenotyping and quantification of cytokine and chemokine protein levels in the lungs. Our previous analysis RNA levels in the lungs of obese/diabetic mice suggested that neutrophils may be present at much higher concentration in obese/diabetic lungs.

As obesity is known to augment the immune system, we established the baseline immune cell population in the lungs of uninfected ND and HFD mice to determine what the initial skew was of inflammatory infiltrates. When we examined the cells of the innate immune system, we observed HFD mice had about 5-fold fewer neutrophils and 3-fold fewer monocyte-derived dendritic cells (moDCs) compared to ND mice (Figure 5A & 5C). For adaptive immune system cells, we observed that HFD mice had around 2-fold higher total T cells, CD4^+^, and CD8^+^ T cells (Figure 6 C-E). With established baselines, we could examine how infection and infection plus vaccination change the landscape of the immune cells.

**Figure 5:**
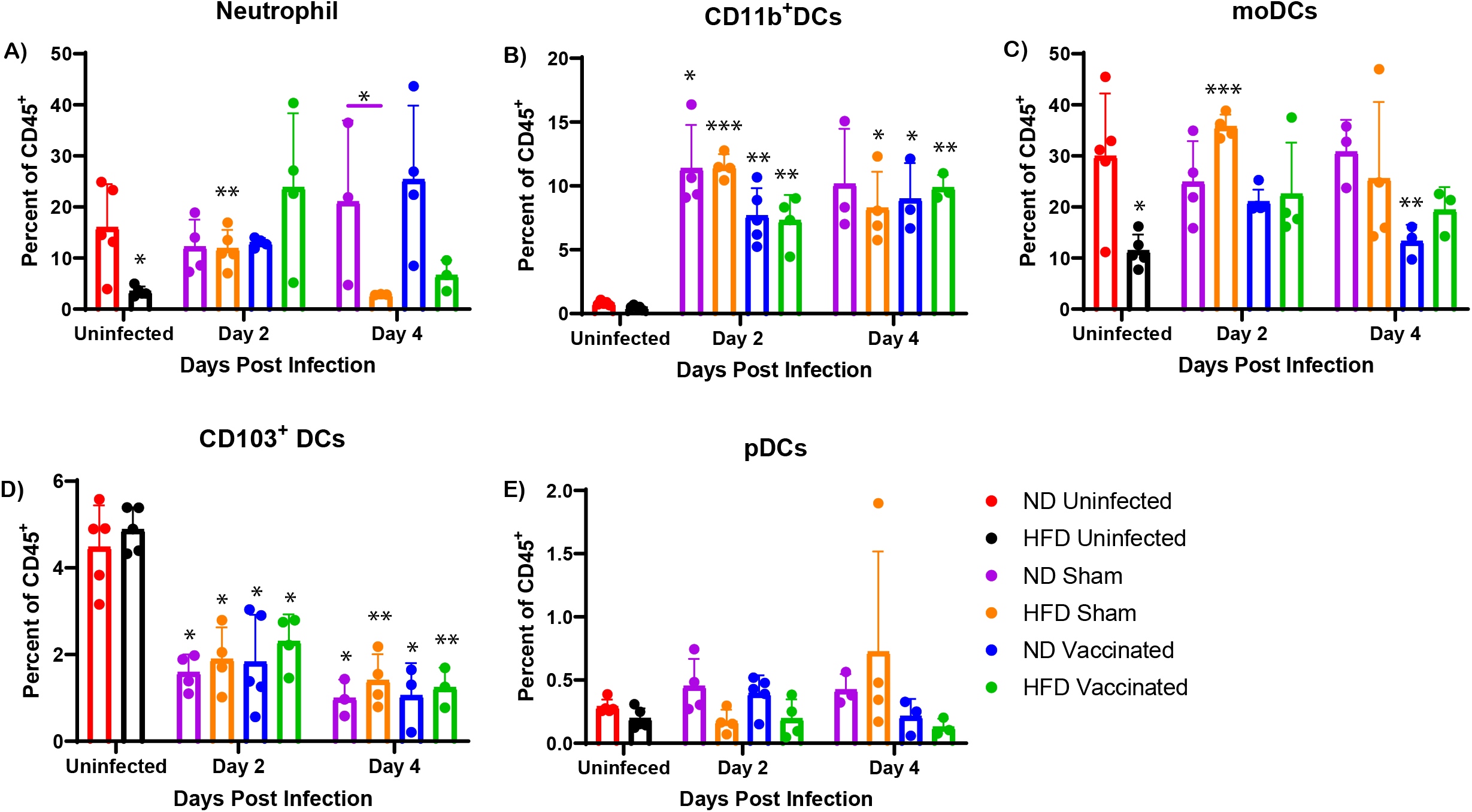
Increased neutrophil recruitment in the lungs of HFD mice after SARS-CoV-2 challenge. After SARS-2 MA10 challenge, lungs were harvested, cells isolated, and stained for surface markers for **A**. neutrophils, **B**. CD11b^+^ DCs, **C**. monocyte-derived DCs, **D**.CD103^+^ cDCs, **E**. plasmacytoid DCs at day 2 and 4 post-challenge. * p < 0.05 and ** p < 0.01 as data were analyzed by mixed-effect analysis followed by Tukey for multiple comparisons *n*=3-5 mice/group.

**Figure 6:**
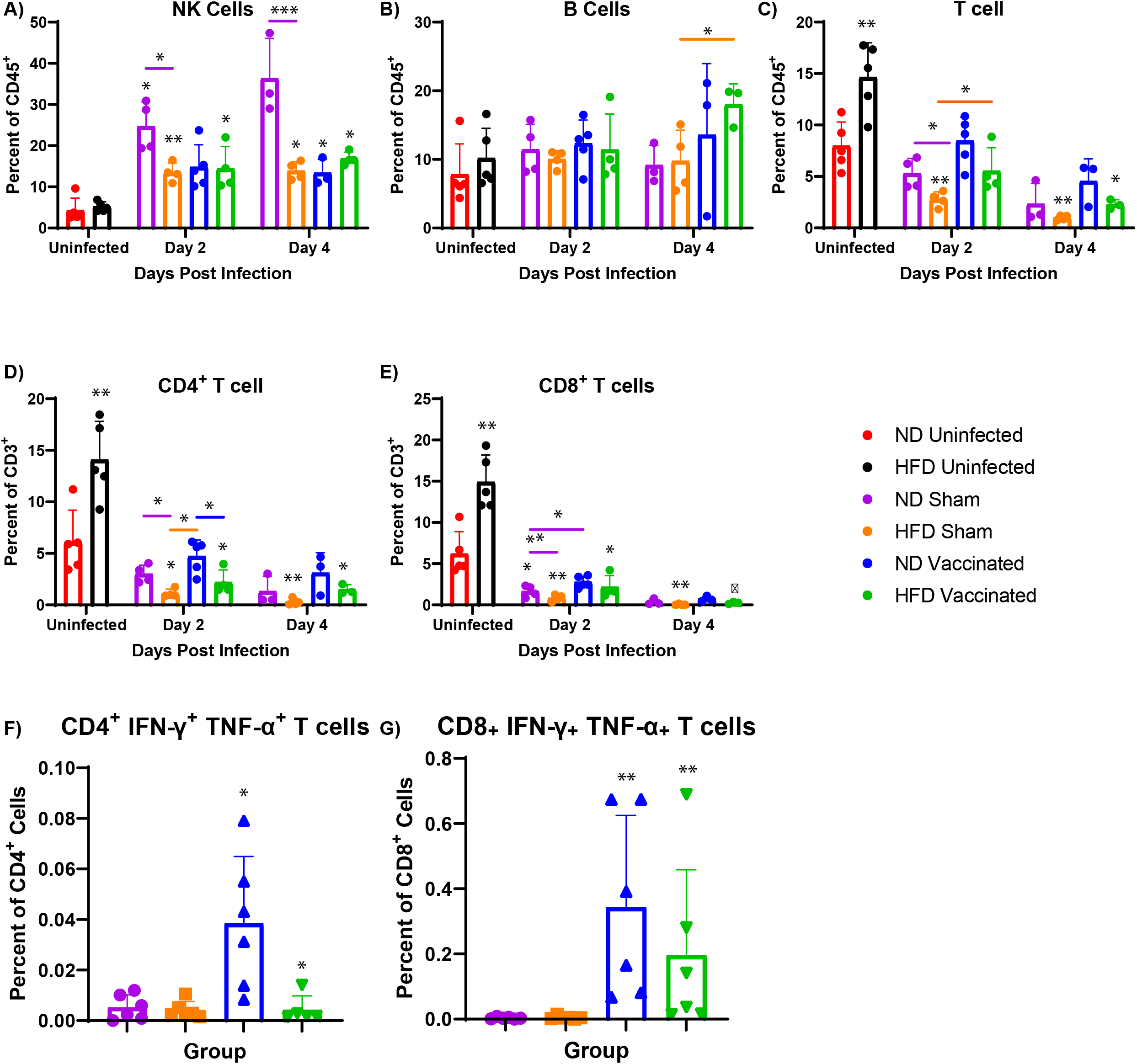
T cell dysregulation in the lungs of HFD mice after SARS-CoV-2 challenge. After SARS-2 MA10 challenge, lungs were harvested, cells isolated, and stained for surface markers for **A**. Natural Killer cells, **B**. B cells, **C**. Total T cells, **D**. T helper cells, **E**. Cytotoxic T cells at day 2 and 4 post-challenge *n*=3-5 mice/group. * p < 0.05 as data were analyzed by mixed-effect analysis followed by Tukey for multiple comparisons *n*=3-5 mice/group **F**. CD4^+^ SARS-CoV-2-specific T cells and **G**. CD8^+^ SARS-CoV-2 specific T cells from naïve and vaccinated ND and HFD mice after *ex-vivo* stimulation *n*=6 mice/group * p <0.5 as determined by an unpaired two-sided t-test. The data are presented as mean ± SD.

As neutrophils are the first cell recruited to the site of infection, we quantified the percentage of neutrophils in the lungs. On day 2 post-infection, unvaccinated HFD mice recruited ∼5-fold the neutrophils to the lungs as compared to uninfected HFD mice (Figure 5A). Interestingly, this recruitment was to the level of unvaccinated ND mice, which did not increase neutrophil recruitment compared to uninfected (Figure 5A). On day 4 post-infection, around a 5-fold decrease in neutrophils was observed between the sham HFD and ND mice, which indicated the HFD mice had returned to baseline (Figure 6A). To determine if other innate immune system cells were augmented during infection, we next examined the different subsets of DCs. Regardless of vaccine status, ND and HFD had ∼20-fold increase in CD11b^+^ DCs and ∼2-fold decrease in CD103^+^ DCs compared to uninfected. We did not observe changes in the moDCs of ND vaccinated or unvaccinated compared to uninfected ND mice (Figure 6C). However, on day 2 post-infection, sham-vaccinated HFD mice had around a 3-fold increase in moDCs compared to uninfected, which was comparable to ND uninfected mice (Figure 5C). On day 4 post-infection, the unvaccinated ND mice had ∼3-fold higher moDCs than vaccinated ND mice. Collectively, this data demonstrates that HFD mice preferentially recruit neutrophils to the site of infection. Their immune system is below the baseline of ND mice and must compensate for this. After examining myeloid lineage cells, we wanted to determine how cells of lymphoid origin responded to infection.

NK cells are lymphoid origin but are cells of the innate immune system and could play a role in the differential protection seen. Unvaccinated ND mice had about a significant 6-fold increase of NK cells at day 2 post-infection, with a trend at day 4 post-infection compared to uninfected (Figure 6A).

Vaccinated ND mice did not increase their percentage of NK cells compared to unvaccinated, which is not surprising as vaccination should elicit a robust adaptive immune response. Regardless of vaccination status, HFD mice had a 2-fold increase of NK cells on day 2 and 4 post infection (Figure 6A). Suggesting that vaccination was not as efficient in these mice. Next, we wanted to examine the cells of the adaptive immune system.

For B cells that produce antibodies, we observed that HFD vaccinated mice had ∼2-fold higher B cells in their lungs at day 4 post-infection than HFD unvaccinated mice (Figure 6B). Finally, we wanted to examine the T cell response between these mouse groups. When we compared uninfected ND and HFD mice, we observed ∼2-fold higher total T cells, CD4^+^ and CD^+^8 T cells (Figure 6C-E). We hypothesized that HFD mice had a lower percentage of innate immune cells which allows for the percentage of T cells to contribute more to the overall CD45^+^ cell population. On day 2 post-infection sham vaccinated ND had ∼2-fold higher T cells, CD4^+^ and CD8^+^ T cells compared to sham HFD mice (Figure 6C-E), and around 2-fold lower CD8^+^ T cell compared to ND vaccinated mice (Figure 6E). Vaccinated ND mice had ∼2-fold CD4^+^ T cells compared to HFD vaccinated mice (Figure 6D). On day 2 post-infection unvaccinated HFD mice had ∼2-fold less total T cells and CD4^+^ T cells when compared to HFD and ND vaccinated mice respectively (Figure 6C&D). This data suggests that HFD mice have a lower percentage of T cells compared to ND mice and have a poorer CD4^+^ T cell response to a challenge after vaccination.

### HFD mice have altered antigen specific T cell activation

We performed *ex vivo* stimulation of T cells with SARS-CoV-2 spike peptide pool from the lungs of vaccinated and unvaccinated ND and HFD mice. For unvaccinated ND and HFD mice, we observed negligible amounts of cytokine production from CD4^+^ or CD8^+^ T cells after stimulation (Figure 6 F&G). It was determined that vaccinated mice production of IFN-γ and TNF-α after peptide stimulation, indicating a T cell response skewed towards a Th_1_ response (Figure 6 F&G). For HFD vaccinated mice, IFN-γ and TNF-α were secreted from CD4^+^ and CD8^+^ T cells, but only CD8^+^ T cells significantly increased compared to unvaccinated (Figure 6 F&G). However, ND vaccinated mice produced IFN-γ and TNF-α from CD4^+^ and CD8^+^ T cells and produced more from CD4^+^ T cells compared to HFD mice (Figure 7 F&G). Taken together, vaccination induces a Th_1_ response in ND and HFD mice, but vaccination does not induce a robust response in HFD mice compared to ND mice which collaborates with could be due to lower percentage of CD4^+^ T cell in HFD mice (Figure 6D).

**Figure 7:**
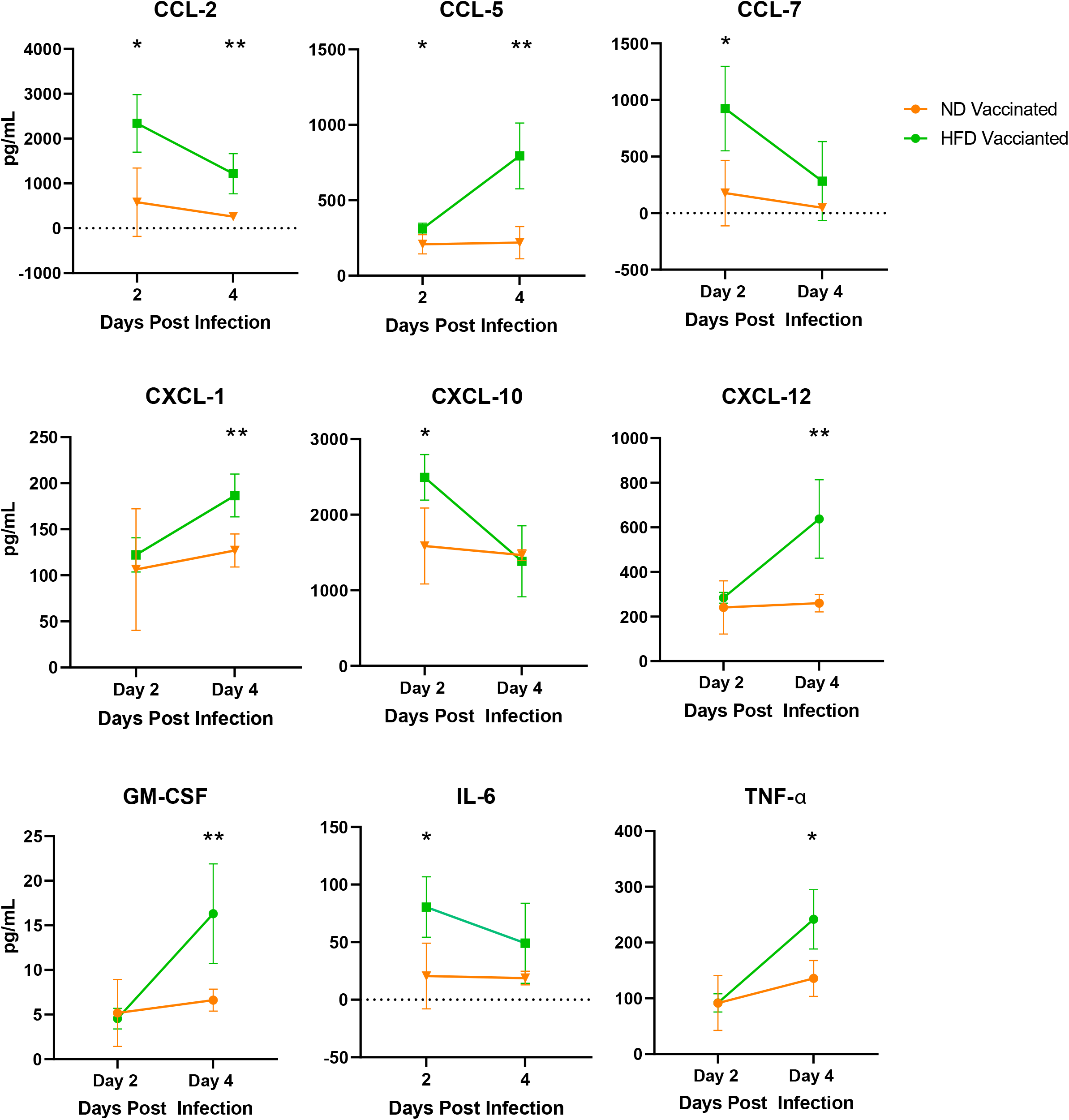
Increased neutrophil recruitment and cytokine production in HFD vaccinated mice after challenge. On day 2,4,7 dpi, lungs were harvested and used in a Bio-plex pro mouse chemokine panel. The pg/mL was generated from a standard curve. * p < 0.05 and ** p < 0.01 as analyzed by mixed-effect analysis followed by Sidak for multiple comparisons *n*=3-5 mice/group.

### HFD Mice have higher levels of cytokine and chemokines in the lungs after challenge

To further characterize the immune responses in ND and HFD mice, we next performed a multiplexing cytokine assay to examine the concentration of soluble immune mediators in the lungs of vaccinated ND and HFD SARS-CoV-2 infected mice. When we compared the ND and HFD mice at 2 dpi, we observed higher levels of CCL2, CCL5, CCL7, CXCL10, and IL-6, and at 4dpi, CCL2, CCL5, CXCL12, CXCL1, GM-CSF, and TNF-α in HFD mice compared to ND mice (Figure 7). These chemokines are all produced from or attract neutrophils and are concordant with the increased neutrophil infiltration we observe in flow cytometry analysis of infected mice. As we observed this infiltration of neutrophils in HFD mice compared to ND mice, we wanted to determine if neutrophils played a role in the increased mortality of HFD mice.

To address the role neutrophils play in the increased mortality of HFD mice, we depleted neutrophils with an anti-Ly6G antibody (clone, 1A8) in the context of a lethal (1×10^5^ pfu/mouse) SARS-CoV-2 mouse model. A day before infection, mice were intra-peritoneally injected with either anti-Ly6G or control (IgG2a isotype) antibody, and administration continued every other day throughout the infection. We examined mortality as our first parameter to determine the effect neutrophils have on increased mortality in HFD mice. ND mice with neutrophils depleted compared to isotype control showed no differences in survivability (Figure 8A). However, HFD mice lacking neutrophils demonstrated increased survivability compared to HFD isotype and anti-Ly6G ND mice (Figure 8A). After determining the effect on mortality, we wanted to examine weight loss. HFD mice given the isotype antibody lost around 10 percent body weight, which was similarly observed in the previous study (Figure 8B & 2C). As previously observed, ND mice lost around 30 percent of their body weight (Figure 8B & 2C). Neutrophil depleted ND mice lost around 20 percent of body weight before death (Figure 8B & 2C). Surprisingly, neutrophil depleted HFD mice lost around 10 percent body weight, but they survived to the experiment’s conclusion, which was not previously observed (Figure 8B & 2C). This data suggests that neutrophil recruitment has a direct role the increased mortality of SARS-CoV-2 infection.

**Figure 8:**
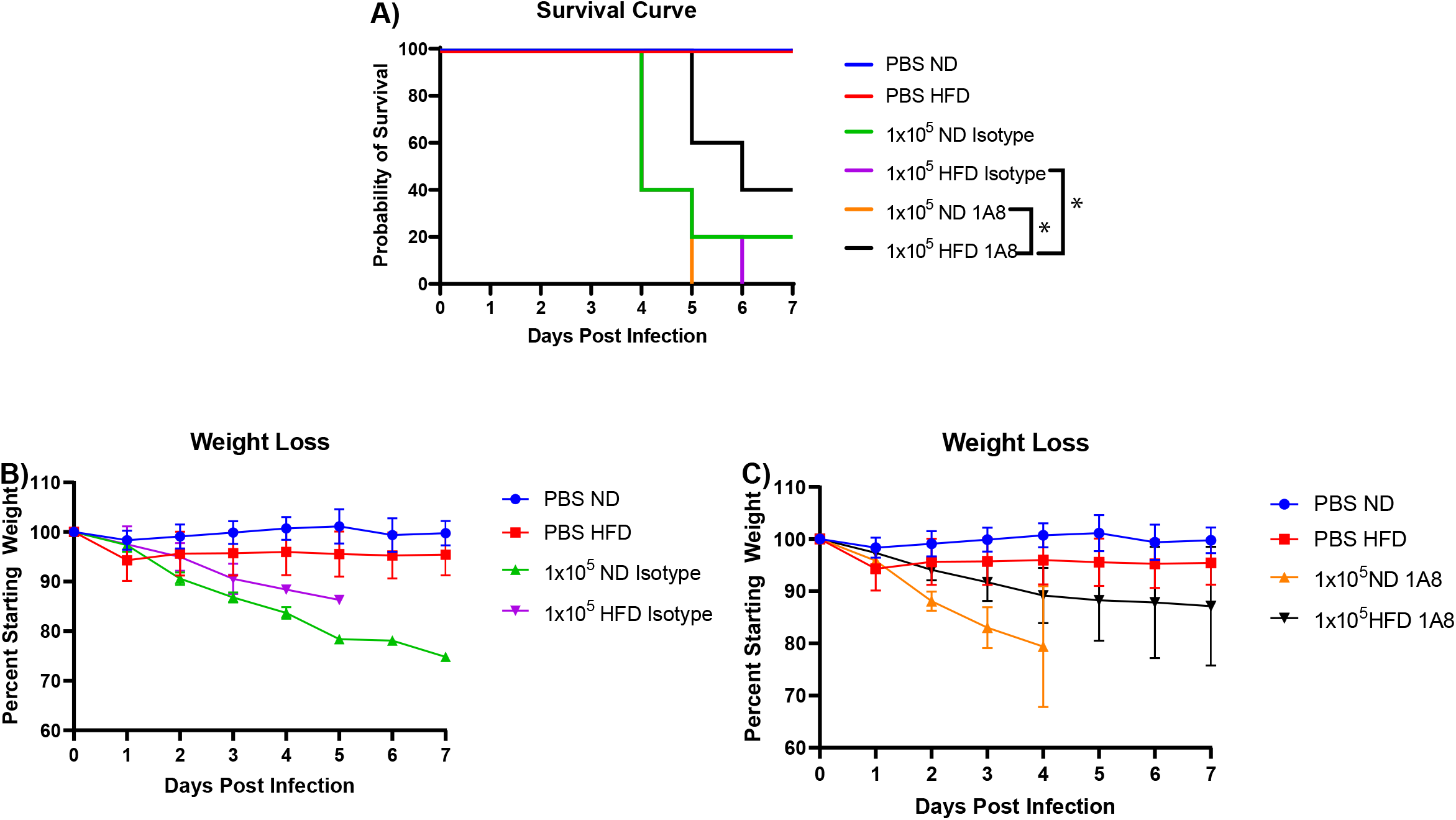
Increased HFD survival with neutrophil depletion after challenge. The **A**. survival and percent weight loss for **B**. isotype antibody control or **C**. anti-Ly6G antibody ND and HFD mice. * p < 0.05 as analyzed by Gehan-Breslow-Wilcoxon test for survival *n*=5 mice/group.

## Discussion

SARS-CoV-2 infection is more severe in obese and diabetic individuals[5-7, 21]. This has been observed in epidemiological data across many countries and economic levels[8]. Increased disease in diabetic and obese individuals is seen following several viral infections, including Influenza virus, Hepatitis B, Hepatitis C, and another highly pathogenic coronavirus, MERS-CoV[19, 41, 42]. We have previously demonstrated that in a mouse model of MERS-CoV, there is increased inflammation and a dysregulated immune response[18]. In this work, we utilized an obese and diabetic mouse model for the study of SARS-CoV-2 pathogenesis to determine whether we could observe increased disease as well as assess the effect of obesity and diabetes on SARS-CoV-2 vaccine responses.

We find that there are significant differences in the obese and diabetic (HFD) mice compared to normal mice after SARS-CoV-2 infection. First, the HFD mice have increased mortality after SARS-CoV-2 infection compared to ND mice. There are no observed differences in lung pathology or virus titer to explain the reason for the mortality other than the increased neutrophil infiltration we observed in the lungs of mice, which may have an overreaching effect on other aspects of host response not accounted for in this work.

Upon vaccination of the ND and HFD mice, we identify severe effects of obesity and diabetes on the vaccine mediated protection in mice. While neutralizing antibody levels are similar between ND and HFD mice vaccinated with NVX-2373 and lethality is protected in both the ND and HFD vaccinated mice, even with their advanced age to the initial testing, there is no protection from weight loss in the vaccinated HFD mice compared to ND mice. When virus levels in the lung were analyzed, we observe a significant delay in clearance of virus at 2 dpi with the HFD mice showing no reduction in virus titer of vaccinated mice while the ND mice reduce virus titer by 4 logs in lungs at the same timepoint. Analysis of the inflammatory response in the mice also shows differences with the key difference being higher levels of neutrophils in the lungs of sham and vaccinated HFD mice compared to ND mice. This was corroborated by assaying for chemokines and cytokines in these lungs. We find that neutrophil produced chemokines and cytokines are significantly upregulated in the HFD mice, potentially associated with the reduction in clearance of virus from HFD vaccinated mice. Analysis of adipose tissue in humans also displays increased inflammation which correlates with what we observe in mice[43, 44].

We interpret these results to demonstrate that the HFD mouse model reflects what is seen in humans with obesity and diabetes throughout the COVID-19 pandemic. This comorbid population has significantly more hospitalizations, irrespective of vaccine status. This model also allows us to begin to differentiate the mechanism behind this lack of protection from SARS-CoV-2. The receptor for SARS-CoV-2 is ACE2 and in humans there is a modest increase in ACE2 levels in some reports suggesting that increased disease could be due to increased entry and thus infection with SARS-CoV-2[45]. In mice, we do not see an increase in ACE2 levels in lungs in the diabetic and obese model used here. While this could be a reason for increased disease in diabetic humans, we do not believe this is the reason for increased disease that we see in mice. With equivalent neutralizing antibody levels but less clearance in lungs, we hypothesize that this is due to changes in quality of the antibodies produced. Potentially the antigens that the antibodies recognize, or the subtypes of the antibodies are changed in the obese/diabetic mice compared to normal mice. There are reports of altered T cell activation in humans with obesity in response to COVID19 vaccination suggesting that T cells may be involved in the altered inflammatory response as well[46]. We also observe the strong association of a hyperactive neutrophil response in the lungs. This could either be due to increased neutrophil recruitment or once there the neutrophils are not responding correctly. We have shown that depleting neutrophils in diabetic/obese mice protects them from lethal disease suggesting neutrophil activity in the lungs is pathologic in this model.

We hypothesize that the dysregulated immune response leads to increased morbidity and mortality as well as skewing the ability of antibodies produced by vaccination to clear viral infection. It is yet to be determined whether it is the metabolic disease associated with obesity that is driving this response or the hyperglycemia of type 2 diabetes however alterations to therapeutic efficacy from vaccination to anti-viral drugs can be affected by either type of metabolic change and will be studied further in the future.

## Methods

### Cells and Virus Strain

Virus and cells were processed as described previously [47]. Briefly, Vero E6 cells (ATCC# CRL 1586) were cultured in DMEM (Quality Biological), supplemented with 10% (v/v) fetal bovine serum (Gibco), 1% (v/v) penicillin/streptomycin (Gemini Bio-products) and 1% (v/v) L-glutamine (2 mM final concentration, Gibco) (Vero media). Cells were maintained at 37°C and 5% CO_2_. SARS-CoV-2 (MA10) was provided by Dr. Ralph Baric (UNC). Stocks were prepared by infection of VeroE6 cells for two days when CPE was starting to be visible [48]. Media were collected and clarified by centrifugation before being aliquoted for storage at −80°C. The stock titer was determined by plaque assay using Vero E6 cells as described previously [49]. All work with infectious virus was performed in a Biosafety Level 3 laboratory and approved by our Institutional Biosafety Committee.

### Animal Ethics Statement

The University of Maryland School of Medicine is accredited by the Association for Assessment and Accreditation of Laboratory Animal Care (AAALAC International). All animal procedures were in accordance with NRC Guide for the Care and Use of Laboratory Animals, the Animal Welfare Act, and the CDC/NIH Biosafety in Microbiological and Biomedical Laboratories. Mouse studies were approved by The University of Maryland School of Medicine IACUC. Studies were conducted in accordance with the National Institutes of Health Guide for Care and Use of Laboratory Animals (NIH publication 8023, Revised 1978). All mouse infections are carried out in an animal BSL-3 facility in accordance with approved practices.

### Husbandry of diet induced obesity mice

18-weeks-old male C57BL/6 diet included obesity (DIO) and C57BL/6 control mice were purchased from Jackson Laboratory. After arrival, mice were kept on a diet of normal chow (22% kcal from fat; Teklad, 2019S) or a diet consisting of high fat chow (60% kcal from fat; Teklad, TD.06414) and maintained on their respective diets throughout the course of the experiments. To evaluate the diabetic status of the mice their fasting blood glucose and starting weight were evaluated before infection. Animals were fasted with water only for six hours prior glucose testing to obtain fasting blood glucose measurements. Sub-mandibular blood was used to obtain blood glucose measurements on a Contour One blood glucose meter strip (Ascensia, Parsippany, NJ).

### Mouse Infections

18-weeks-old male C57BL/6 diet included obesity (DIO) and C57BL/6 control mice were purchased from Jackson Laboratory. On day 0, mice were anesthetized by intraperitoneal injection 50 μL of a mix of xylazine (0.38 mg/mouse) and ketamine (1.3 mg/mouse) diluted in phosphate-buffered saline (PBS).

While anesthetized, mice were intranasally inoculated with either 50 µL sterile PBS, 1 × 10^3^ PFU or 1×10^5^ PFU of SARS-CoV-2 MA10. Mice are monitored daily for weight loss and signs of morbidity. On days 2, 4, and 7, animals were euthanized, and lung tissue was collected for further analysis.

### Mouse Challenge Study

18-weeks-old male C57BL/6 diet included obesity (DIO) and C57BL/6 control mice were purchased from Jackson Laboratory and immunized by intramuscular (IM) injection with two doses spaced 14 days apart (day 0 and 14) of 1 µg rS-WU1, with 5 μg saponin-based Matrix-M™ adjuvant (Novavax, AB, Uppsala, SE) as a prime/boost. A placebo group was injected with vaccine formulation buffer as a negative control.

Serum was collected for analysis on days 0 and 28. Vaccinated and control animals were intranasally challenged with SARS-CoV-2 on study day 32. On days 2 and 4 post-challenge, animals were euthanized, and lung tissue was collected for further analysis.

### Histology & Immunohistochemistry

Lung sections were fixed in 4% paraformaldehyde (PFA) in phosphate-buffered saline (PBS) for a minimum of 48 h, after which they were sent to the Histology Core at the University of Maryland, Baltimore, for paraffin embedding and sectioning. Five-micrometer sections were prepared and used for hematoxylin and eosin (H&E) staining by the Histology Core Services.

For SARS-CoV-2 nucleocapsid immunohistochemistry staining, five-micrometer sections were cut from paraffin-embedded blocks and placed onto positively charged slides. The sections were immunostained using the Dako EnVision FLEX + detection system (DAKO, Carpinteria, CA). The Dako PT link was used for deparaffinization and heat-induced epitope retrieval using Dako Target Retrieval Solution, High pH, for 20 minutes. Endogenous peroxidase activity was blocked with DAKO Peroxidase-Blocking Reagent for 5 min before incubation with rabbit polyclonal SARS Nucleocapsid Protein Antibody [NB100-56576]

(Novus Biologicals, Centennial, CO), 1:400 for 20 minutes at room temperature, followed by DAKO Anti-rabbit HRP Detection Reagent for 20 minutes. Finally, the sections were incubated for 10 min with DAKO diaminobenzene (DAB) before they were counterstained with Dako FLEX hematoxylin, rinsed, and mounted in Cytoseal XYL (Thermo Scientific, Waltham, MA). Sections were imaged at 10x magnification, and figures were put together using Adobe Photoshop and Illustrator software (Adobe, San Jose, CA).

### Plaque Assay

VeroE6/TMPRSS2 cells were cultured in DMEM (Quality Biological), supplemented with 10% (v/v) heat-inactivated fetal bovine serum (Sigma), 1% (v/v) penicillin/streptomycin (Gemini Bio-products), and 1% (v/v) L-glutamine (2 mM final concentration, Gibco) (Vero Media). Cells were maintained at 37°C (5% CO_2_). SARS-CoV-2 lung titers were quantified by homogenizing mouse lungs in 1ml phosphate-buffered saline (PBS; Quality Biological Inc.) using 1.0mm glass beads (Sigma Aldrich) and a Beadruptor (Omni International Inc). VeroE6 cells are plated in 12 well plates with 1.5×10^5^ cells per well. SARS-CoV-2 virus titer in plaque forming units was determined by plaque assay. In the plaque assay, 25 µl of the lung homogenate is added to 225 µl of DMEM and diluted 10-fold across a 6-point dilution curve with 200ul of diluent added to each well. After 1 hour, a 2 ml agar overlay containing DMEM is added to each well. Plates are incubated for three days at 37°C (5% CO_2_) before plaques are counted.

### SARS-CoV-2 Neutralization Titer Determination

SARS-CoV-2 neutralizing antibody titers were determined as described previously [50]. Briefly, serum samples were heat-inactivated to remove complement and allowed to equilibrate to room temperature before processing. Samples were diluted in duplicate for a 12-point, 1:2 dilution series with an initial dilution of 1:40 dilution (1:80 following virus addition). Dilution plates were then transported into the BSL-3 laboratory, and equal volume SARS-CoV-2 inoculum was added to each plate. A non-treated, virus-only control and a mock infection control were included on every plate. The sample/virus mixture was then incubated at 37°C (5.0% CO_2_) for 1 hour before transferring to 96-well titer plates with confluent VeroE6 cells. Titer plates were incubated at 37°C (5.0% CO_2_) for 72 hours, followed by CPE determination for each well in the plate. The first sample dilution to show CPE was reported as the minimum sample dilution required to neutralize >99% of the concentration of SARS-CoV-2 tested (neut99).

### Gene Expression

Mouse lung tissue was homogenized in 1 mL of TRIzol (Ambion) using 1.4-mm glass beads (Omni International, Inc.) and a beadruptor (Omni International, Inc.) RNA was extracted using the DIRECT-zol Mini RNA Extraction kit (Zymo Research) per the manufacturer’s instructions. RNA was transcribed into cDNA using RT^2^ First Strand Kit (Qiagen). Gene expression was performed using Real-time PCR for RT^2^ Profile PCR Array Mouse Cytokine and Chemokines using RT^2^ SYBR® Green Mastermixes (Qiagen). Using the RT^2^ qPCR Array Data Analysis spreadsheet, gene expression was calculated using the ΔΔCt method with housekeeping genes as the control, and fold-change calculated relative to control mice infected lungs. Fold-regulation, which represents fold-change in a biologically meaningful way, was calculated using fold change. A fold-change value greater than one indicates an up-regulation, and the fold-regulation is equal to the fold change. A fold-change value less than one indicates a down-regulation, and the fold-regulation is negative inverse of the fold-change.

### Cytokine and Chemokine Quantification

For the quantification of secreted cytokine and chemokines induced by SARS-CoV-2 infection, mice lung tissue was homogenized in 1 mL of PBS (Quality Biologicals, Inc.) using 1.4-mm glass beads (Omni International, Inc.) and a beadruptor (Omni International, Inc.). Bio-Plex Pro Mouse Chemokine Panel 31-plex (BIORAD) was used to quantify cytokines and chemokines in lung homogenate according to the manufacturer’s protocol with the addition of a fixation step after the final wash. Samples underwent a 24-hour fixation (4 % formaldehyde, Sigma-Aldrich) at 4°C. After fixation, the plate was washed three times in wash buffer, microparticles were resuspended in 100 µL wash buffer, and analyzed on a Luminex MagPix and xPONENT Software version 4.3. Based on the standard curve, concentrations (pg/mL) of each analyte were calculated for all samples.

### Flow Cytometry

Single cells were isolated from lungs using the Lung Dissociation Kit, mouse and gentleMACS Dissociator (Miltenyi Biotec). Lungs were placed in C tubes (Miltenyi Biotec) containing 2.2 mL Buffer S, 15 µL of Enzyme A, and 100 µL of Enzyme D. Lungs were run on m_lung_01 and allowed to digest for 30 minutes at 37°C followed by running m_lung_02. Cell suspensions were filtered on a 70-μm filter (BD), and cells were pelleted by centrifugation. Red blood cells were lysed in ammonium-chloride-potassium lysis buffer (Quality Biological, Inc.) and subsequently washed with PBS containing 3% FBS. Approximately, 1×10^6^ cells were plated and washed twice with PBS containing. Cells were stained for viability using the Live/Dead Fixable NIR Dead Cell Stain Kit (Molecular Probes). Cells were washed with PBS containing 3% FBS. Cells were stained with antibody cocktails made in BD Horizon Brilliant Stain Buffer (BD). The antibodies used were as follows: CD45 Alexa Fluor 700 (BioLegend, clone 30-F11), CD11b Brilliant Violet 650 (BioLegend, clone M1/70), Ly6G Brilliant Violet 785 (BioLegend, clone 1A8), Ly6C Brilliant Violet 510 (BioLegend, clone HK1.4), CD3 BUV 563 (BD, clone 145-2C11), CD4 FITC (BioLegend, clone GK1.5), CD8a APC (BioLegend clone 53-6.7), CD19 PE (BioLegend clone 6D5), and NK1.1 Pacific Blue (BioLegend, clone PK136), CD65 Brilliant Violet 605 (BioLegend, clone X54-5/7.1), CD103 BUV 661 (BD, clone 2e7), Siglec-H BUV 563 (BD, clone 440c), and Siglec-F Brilliant Violet 711 (BD, clone E50-2440).

Stained cells were washed twice and fixed for 1 hour in FluoroFix (BioLegend). For Intracellular Cytokine Staining, lung leukocytes 2×10^6^ cells/well from vaccinated and sham mice were cultured at 37°C for 5 hours in the presence of SARS-CoV-2 spike peptide pool (1µg/mL) kindly provided by Alessandro Sette LA Jolla Institute, Brefeldin A and Monensin (BioLegend). The peptide pool consisted of overlapping 15-mers by 10 spanning entire protein. The treated cells were incubated with LIVE/DEAD Fixable NIR Dead Cell stain (Invitrogen) for 20 min at room temperature. Cells were then labeled for cell surface markers at 4°C for 15 minutes in the dark followed by fixation and permeabilization for 20 min at 4°C (BD CytoFix/CytoPerm) then labeled with intracellular antibody cocktail. The antibodies used were as follows: IFN-γ BV421 (BioLegend, clone: XMG1.2), TNF-α BV650 (BioLegend, clone:MP6-XT22), IL-4 BV605 (BioLegend, clone: 11B11), and IL-17a BV711 (BioLegend, clone: TC11-18H10.1). Fixed cells were washed once in PBS containing 3% FBS and resuspended in PBS containing 3% FBS. Samples were acquired (approximately 50,000 events) using the Aurora-UV spectral flow cytometer (Cytek), and data were analyzed using FCS Express analysis software (DeNova Software, Pasadena, CA).

### Neutrophil Depletion

18-weeks-old male C57BL/6 diet included obesity (DIO) and C57BL/6 control mice were purchased from Jackson Laboratory and neutrophils were depleted with 500 µg of anti-Ly6G (clone 1A8) or isotype antibody (IgG2a isotype) (Bio-X-Cell, Lebanon, NH), via i.p. injection 1 day prior to infection with SARS-CoV-2 and every two days afterwards. Mice were infected with either 50 µL sterile PBS or 1×10^5^ PFU of SARS-CoV-2 MA10. Mice are monitored daily for weight loss and signs of morbidity. On day 7, animals were euthanized, and lung tissue was collected for further analysis.

### Statistical Analysis

Statistical analyses were performed with GraphPad Prism 9.2 software (GraphPad Software, San Diego, CA). Data were analyzed using unpaired t-test, Log-rank (Mantel-Cox) test, one-way or two-way ANOVA followed by Tukey, Sidak, or Dunnett multiple comparison post-hoc test as indicated. All statistical analysis were two-sided and a p < 0.05 was consisted as statistically significant.

## References

1. CDC. National Diabetes Statistics Report. Available from: https://www.cdc.gov/diabetes/data/statistics-report/index.html.

2. Esser, N., et al., Inflammation as a link between obesity, metabolic syndrome and type 2 diabetes. Diabetes Res Clin Pract, 2014. 105(2): p. 141–50.

3. Ritchie, S.A. and J.M. Connell, The link between abdominal obesity, metabolic syndrome and cardiovascular disease. Nutr Metab Cardiovasc Dis, 2007. 17(4): p. 319–26.

4. Grundy, S.M., Obesity, metabolic syndrome, and cardiovascular disease. J Clin Endocrinol Metab, 2004. 89(6): p. 2595–600.

5. Kompaniyets, L., et al., Body Mass Index and Risk for COVID-19-Related Hospitalization, Intensive Care Unit Admission, Invasive Mechanical Ventilation, and Death - United States, March-December 2020. MMWR Morb Mortal Wkly Rep, 2021. 70(10): p. 355–361.

6. Apicella, M., et al., COVID-19 in people with diabetes: understanding the reasons for worse outcomes. Lancet Diabetes Endocrinol, 2020. 8(9): p. 782–792.

7. Rawshani, A., et al., Severe COVID-19 in people with type 1 and type 2 diabetes in Sweden: A nationwide retrospective cohort study. Lancet Reg Health Eur, 2021. 4: p. 100105.

8. Ozonoff, A., et al., Phenotypes of disease severity in a cohort of hospitalized COVID-19 patients: Results from the IMPACC study. EBioMedicine, 2022. 83: p. 104208.

9. Heubner, L., et al., Extreme obesity is a strong predictor for in-hospital mortality and the prevalence of long-COVID in severe COVID-19 patients with acute respiratory distress syndrome. Sci Rep, 2022. 12(1): p. 18418.

10. Rosero, P.A., et al., Risk Factors for COVID-19: A Systematic Mapping Study. Stud Health Technol Inform, 2022. 299: p. 63–74.

11. Badawi, A. and S.G. Ryoo, Prevalence of Diabetes in the 2009 Influenza A (H1N1) and the Middle East Respiratory Syndrome Coronavirus: A Systematic Review and Meta-Analysis. J Public Health Res, 2016. 5(3): p. 733.

12. Rathnasinghe, R., et al., The N501Y mutation in SARS-CoV-2 spike leads to morbidity in obese and aged mice and is neutralized by convalescent and post-vaccination human sera. medRxiv, 2021.

13. Johnson, R.M., et al., Diet Induced Obesity and Diabetes Enhance Mortality and Reduces Vaccine Efficacy for SARS-CoV-2. bioRxiv, 2022.

14. Assiri, A., et al., Epidemiological, demographic, and clinical characteristics of 47 cases of Middle East respiratory syndrome coronavirus disease from Saudi Arabia: a descriptive study. Lancet Infect Dis, 2013. 13(9): p. 752–61.

15. Cha, R.H., et al., Renal Complications and Their Prognosis in Korean Patients with Middle East Respiratory Syndrome-Coronavirus from the Central MERS-CoV Designated Hospital. J Korean Med Sci, 2015. 30(12): p. 1807–14.

16. Choi, W.S., et al., Clinical Presentation and Outcomes of Middle East Respiratory Syndrome in the Republic of Korea. Infect Chemother, 2016. 48(2): p. 118–26.

17. Gao, Y.D., et al., Risk factors for severe and critically ill COVID-19 patients: A review. Allergy, 2021. 76(2): p. 428–455.

18. Kulcsar, K.A., et al., Comorbid diabetes results in immune dysregulation and enhanced disease severity following MERS-CoV infection. JCI Insight, 2019. 4(20).

19. Yang, Y.M., et al., Impact of Comorbidity on Fatality Rate of Patients with Middle East Respiratory Syndrome. Sci Rep, 2017. 7(1): p. 11307.

20. Rydell-Tormanen, K. and J.R. Johnson, The Applicability of Mouse Models to the Study of Human Disease. Methods Mol Biol, 2019. 1940: p. 3–22.

21. Alanazi, K.H., et al., Diabetes Mellitus, Hypertension, and Death among 32 Patients with MERS-CoV Infection, Saudi Arabia. Emerg Infect Dis, 2020. 26(1): p. 166–168.

22. Lee, F.Y., et al., Phenotypic abnormalities in macrophages from leptin-deficient, obese mice. Am J Physiol, 1999. 276(2): p. C386–94.

23. Manicone, A.M., et al., Diet-induced obesity alters myeloid cell populations in naive and injured lung. Respir Res, 2016. 17: p. 24.

24. O’Brien, K.B., et al., Impaired wound healing predisposes obese mice to severe influenza virus infection. J Infect Dis, 2012. 205(2): p. 252–61.

25. Milner, J.J., et al., Diet-induced obese mice exhibit altered heterologous immunity during a secondary 2009 pandemic H1N1 infection. J Immunol, 2013. 191(5): p. 2474–85.

26. Bandaru, P., H. Rajkumar, and G. Nappanveettil, Altered or Impaired Immune Response to Hepatitis B Vaccine in WNIN/GR-Ob Rat: An Obese Rat Model with Impaired Glucose Tolerance. ISRN Endocrinol, 2011. 2011: p. 980105.

27. Guebre-Xabier, M., et al., NVX-CoV2373 vaccine protects cynomolgus macaque upper and lower airways against SARS-CoV-2 challenge. Vaccine, 2020. 38(50): p. 7892–7896.

28. Tian, J.H., et al., SARS-CoV-2 spike glycoprotein vaccine candidate NVX-CoV2373 immunogenicity in baboons and protection in mice. Nat Commun, 2021. 12(1): p. 372.

29. Leist, S.R., et al., A Mouse-Adapted SARS-CoV-2 Induces Acute Lung Injury and Mortality in Standard Laboratory Mice. Cell, 2020. 183(4): p. 1070–1085 e12.

30. Paich, H.A., et al., Overweight and obese adult humans have a defective cellular immune response to pandemic H1N1 influenza A virus. Obesity (Silver Spring), 2013. 21(11): p. 2377–86.

31. Kim, Y.H., et al., Diet-induced obesity dramatically reduces the efficacy of a 2009 pandemic H1N1 vaccine in a mouse model. J Infect Dis, 2012. 205(2): p. 244–51.

32. Costanzo, A.E., et al., Obesity impairs gammadelta T cell homeostasis and antiviral function in humans. PLoS One, 2015. 10(3): p. e0120918.

33. Fisher-Hoch, S.P., C.E. Mathews, and J.B. McCormick, Obesity, diabetes and pneumonia: the menacing interface of non-communicable and infectious diseases. Trop Med Int Health, 2013. 18(12): p. 1510–9.

34. Hagau, N., et al., Clinical aspects and cytokine response in severe H1N1 influenza A virus infection. Crit Care, 2010. 14(6): p. R203.

35. Painter, S.D., I.G. Ovsyannikova, and G.A. Poland, The weight of obesity on the human immune response to vaccination. Vaccine, 2015. 33(36): p. 4422–9.

36. Tagliabue, C., et al., Obesity: impact of infections and response to vaccines. Eur J Clin Microbiol Infect Dis, 2016. 35(3): p. 325–31.

37. Caimi, B., et al., Sero-survey on long-term care facility residents reveals increased risk of sub-optimal antibody response to BNT162b2: implications for breakthrough prevention. BMC Geriatr, 2022. 22(1): p. 191.

38. Soetedjo, N.N.M., et al., Antibody response following SARS-CoV-2 vaccination among patients with type 2 diabetes mellitus: A systematic review. Diabetes Metab Syndr, 2022. 16(2): p. 102406.

39. Anand, S., et al., SARS-CoV-2 Vaccine Antibody Response and Breakthrough Infection in Patients Receiving Dialysis. Ann Intern Med, 2022. 175(3): p. 371–378.

40. Mitsunaga, T., et al., The evaluation of factors affecting antibody response after administration of the BNT162b2 vaccine: a prospective study in Japan. PeerJ, 2021. 9: p. e12316.

41. Almond, M.H., et al., Obesity and susceptibility to severe outcomes following respiratory viral infection. Thorax, 2013. 68(7): p. 684–6.

42. Honce, R. and S. Schultz-Cherry, Impact of Obesity on Influenza A Virus Pathogenesis, Immune Response, and Evolution. Front Immunol, 2019. 10: p. 1071.

43. Stefan, N., SARS-CoV-2 fires up inflammation in adipose tissue. Nat Rev Endocrinol, 2023. 19(1): p. 8–9.

44. El-Sayed Moustafa, J.S., et al., ACE2 expression in adipose tissue is associated with cardio-metabolic risk factors and cell type composition-implications for COVID-19. Int J Obes (Lond), 2022. 46(8): p. 1478–1486.

45. Wijnant, S.R.A., et al., Expression of ACE2, the SARS-CoV-2 Receptor, in Lung Tissue of Patients With Type 2 Diabetes. Diabetes, 2020. 69(12): p. 2691–2699.

46. Kelly, N.E.W., et al., Antigen specific T cells in people with obesity at five months following ChAdOx1 COVID-19 vaccination. Int J Obes (Lond), 2023. 47(1): p. 83–86.

47. Weston, S., et al., Broad Anti-coronavirus Activity of Food and Drug Administration-Approved Drugs against SARS-CoV-2 In Vitro and SARS-CoV In Vivo. J Virol, 2020. 94(21).

48. Matsuyama, S., et al., Enhanced isolation of SARS-CoV-2 by TMPRSS2-expressing cells. Proc Natl Acad Sci U S A, 2020. 117(13): p. 7001–7003.

49. Coleman, C.M. and M.B. Frieman, Growth and quantification of MERS-CoV infection. Current Protocols in Microbiology, 2015. 37(15): p. 2–15.

50. Keech, C., et al., Phase 1-2 Trial of a SARS-CoV-2 Recombinant Spike Protein Nanoparticle Vaccine. N Engl J Med, 2020. 383(24): p. 2320–2332.

